# Decoding pH-Driven Phase Transition of Lipid Nanoparticles

**DOI:** 10.1101/2024.11.27.625717

**Authors:** Marius F.W. Trollmann, Rainer A. Böckmann

## Abstract

The functionality of lipid nanoparticles (LNPs) as delivery systems in mRNA-based therapeutics is intricately linked to the protonation behavior of their aminolipid components. This study employs large-scale constant-pH molecular dynamics (CpHMD) simulations to decode the environment-dependent pK_a_ of aminolipids in the *Comirnaty* lipid formulation, providing a detailed view of their pH-dependent structural dynamics. Our results reveal a significant shift in the apparent pK_a_ of the aminolipid ALC-0315, from an intrinsic value of 9.3 in water to 4.9 within the LNP environment. This shift arises from the interplay between lipid reorganization and local electrostatic interactions, resulting in distinct protonation states across the LNP core and surface.

At low pH, protonated aminolipids dominate the LNP surface, promoting efficient mRNA encapsulation, whereas at neutral pH, deprotonated aminolipids migrate to the hydrophobic core, driving structural stabilization. Notably, the localized pK_a_ of aminolipids varies significantly with their position, decreasing from near-surface regions (7 to 8) to the hydrophobic core (≤4). These findings elucidate the molecular mechanisms underpinning LNP phase transitions and highlight the key role of pK_a_ shifts for the design of aminolipids and for optimizing LNP compositions for enhanced therapeutic delivery. This study bridges experimental observations with molecular-level insights, advancing the rational development of next-generation lipid-based nanocarriers.

## Introduction

Lipid nanoparticles (LNPs) have become a well-established platform for delivering polynucleotides such as mRNA and siRNA,^1^ as demonstrated by therapeutic products such as Onpattro^2^ and SARS-CoV-2 vaccines.^3–5^ This delivery technology holds significant promise for future vaccines targeting cancers^6,7^ and infectious diseases, e.g., malaria.^8,9^ For instance, BioNTech has developed three mRNA-based vaccines (BNT142, BNT152, and BNT153), all currently in phase 1 or phase 2 clinical trials (NCT05262530, NCT04710043) for treating solid tumors. Another LNP-based vaccine, NBF-006 from Nitto BioPharma, has already completed Phase 1 (NCT03819387) for non-small cell lung, pancreatic, and colorectal cancers.

In LNP formulations, the lipid components play a crucial role in determining the immunogenicity and therapeutic efficacy of the delivered nucleotides.^10^ Most LNPs consist of four key components:^2,11^ (1) ionizable aminolipids (AL), capable of changing their protonation state under physiological conditions;^12^ (2) PEGylated lipids, which control particle size, ^13,14^ prevent aggregation,^14,15^ and enhance circulation time in the body after administration; ^16,17^ (3) cholesterol; and (4) helper phospholipids like 1,2-Distearoyl-sn-glycero-3-phosphocholine (DSPC), which improve stability and transfection efficiency of the LNPs.^14,18^ Together, these lipids encapsulate and protect mRNA from environmental degradation,^11,19^ enabling enhanced transfection compared to unencapsulated mRNA.^10^

Two key properties of ionizable aminolipids influencing transfection efficiency are their pK_*a*_ and molecular shape: While the pK_*a*_ of aminolipids typically ranges between 8.5 and 9.5^12^ and thus outside the physiological range, lipid nanoparticles display pK_*a*_ values from 6 to 7, thus aligning with the acidic environments of the early (pH 6.5) and late endosomes (pH 5.5).^20^ Protonation enhances interactions of LNPs with the negatively charged endosomal membrane, leading to membrane disruption and facilitating the release of LNP cargo, potentially via the formation of an H_II_-phase.^21,22^ This process is supported by the conical molecular shapes of aminolipids like ALC-0315 ([(4-Hydroxybutyl)imino]di-6,1-hexanediyl bis(2-hexyldecanoate)) used in Pfizer-BioNTech’s *Comirnaty*.^3^

Although a variety of experimental^12,23–29^ and simulation-based^19,26,28,30–37^ methods have advanced our understanding of LNP size, structure, lipid distribution within LNP core and shell regions, or the fusion of LNPs with membranes, the shift in the protonation equilibrium — from aminolipids 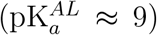 to LNPs 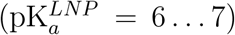 — and the associated pH-dependent charge distributions within LNPs, which are critical for cargo release, remain poorly understood.^38^

In this study, we investigate the protonation behavior and charge distribution of amino-lipids in LNP-mimetic systems at atomistic resolution, employing a constant-pH molecular dynamics (CpHMD) approach based on *λ*-dynamics.^39,40^ This approach allows the study of hundreds of titratable sites on the microsecond timescale. We establish a framework for studying the spatiotemporal protonation dynamics of aminolipids within LNP-like systems at varying environmental pH. Specifically, we parameterized the aminolipid ALC-0315, and based on extensive simulations report the *pH-dependent* structure and composition of *Comirnaty* LNP-mimetic systems, both in presence and in absence of mRNA. Our findings highlight a notable, position-dependent shift in the 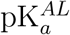 of ALC-0315 when incorporated into a realistic LNP-lipid environment, that rationalizes the experimentally reported 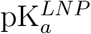 of Comirnaty LNPs. At neutral pH, a significant fraction of protonated aminolipids was observed within the LNP surface layer, emphasizing their critical role in the biophysical properties and functionality of the nanoparticle.

## Results and Discussion

The pH-dependent structural organization of LNP-mimetic systems, LNP core and shell lipid compositions, and the distribution of protonated and deprotonated aminolipids, along with their related pK_a_ values, were analyzed from unbiased microsecond all-atom constant-pH MD (CpHMD) simulations of LNP-mimetic membrane patches reflecting the lipid composition of *Comirnaty* LNPs (systems D, E, F, see Tab. 1 and Fig. 1a). The initial configuration for the CpHMD simulations at different pH values was a lipid bilayer (low pH state, see Fig. 1a). This phase was previously obtained in classical MD simulations with all aminolipids in their protonated state, by studying the spontaneous self-aggregation of lipids starting from a random mixed state.^19^ It is important to note that artificial restraints on the simulation system should be avoided as they may impede pH-dependent LNP phase transitions.^41,42^

**Figure 1:**
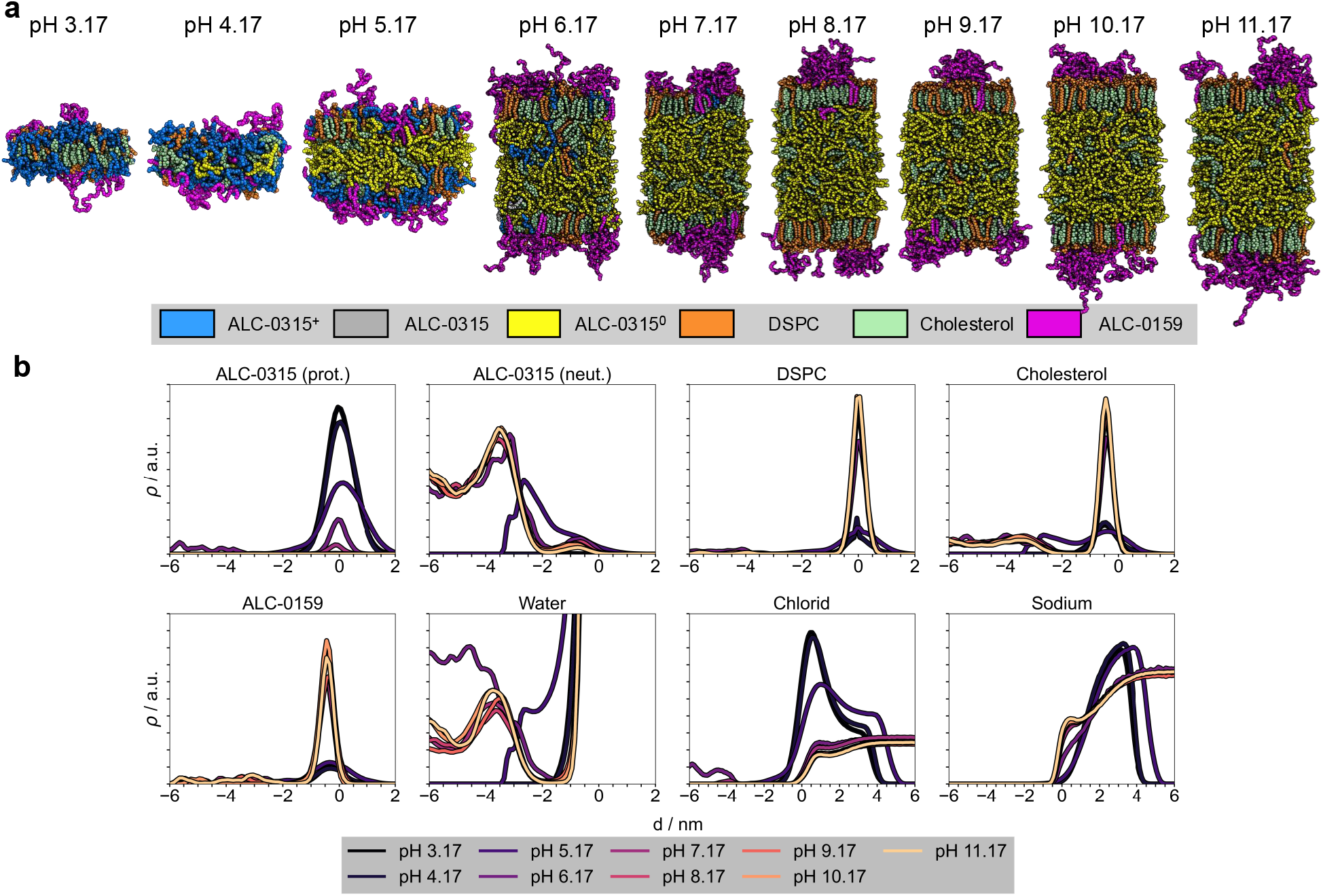
pH-dependent LNP phases. **a** Snapshots of the LNP composition at different pH values. As the pH increases, the aminolipids transition from a protonated state (*blue*) to a deprotonated state (*yellow*), forming a hydrophobic core together with cholesterol (*pale-green*). For pH 5-11, the LNP-mimetic patches were quadruplicated with respect to the initial low-pH structures (pH 3, pH 4) to allow for large-scale phase transitions. DSPC (*orange*), ALC-0159 (*magenta*), and the remaining cholesterol (*green*) accumulate at the boundaries with the solvent phase. ALC-0315 without a specific protonation state (0.2 ≤ *λ* ≤ 0.8) is shown in *gray*. Images were rendered using PyMOL^43^ **b** Average mass densities for all molecule types relative to the membrane surface, defined by the median *z*-position of the phosphorus atoms in DSPC in one leaflet, at different pH levels. The density curves were calculated using the following atoms for each molecule type: nitrogen for ALC-0315, phosphorus for DSPC, oxygen for cholesterol, nitrogen for ALC-0159, and oxygen for water. Density curves represent averages taken after equilibration of *S*^deprot^ values (Supplementary Fig. 6), and were scaled to facilitate comparison between molecule types.

### pH-driven phase transition in *Comirnaty* LNP formulation

The structural and phase behavior of the *Comirnaty* LNP formulation was investigated across an environmental pH range of 3 to 11. At pH 3, almost all aminolipids are in their protonated state, and at pH ≥ 7 transitioned to the deprotonated state (Fig. 1a, protonated aminolipids in *blue*, deprotonated in *yellow*). This shift profoundly influenced the molecular organization of the lipid membrane. At pH 3 and 4, the bilayer structure remained stable throughout microsecond-long CpHMD simulations, consistent with previous findings using fully protonated aminolipids in classical MD. ^19^ Differently, for pH values ≥ 7, the bilayer underwent a phase transition, stabilizing after 9 … 12 *µ*s of all-atom simulation into a LNP-mimetic phase (Supplementary Figures 5–7). This phase is characterized by a hydrophobic core composed of ≈ 70% aminolipids and ≈ 30% cholesterol (Fig. 2), flanked by lipid monolayers. For intermediate pH values between 5 and 6, the system is observed to be in a slowly converging, intermediate phase (see also below).

**Figure 2:**
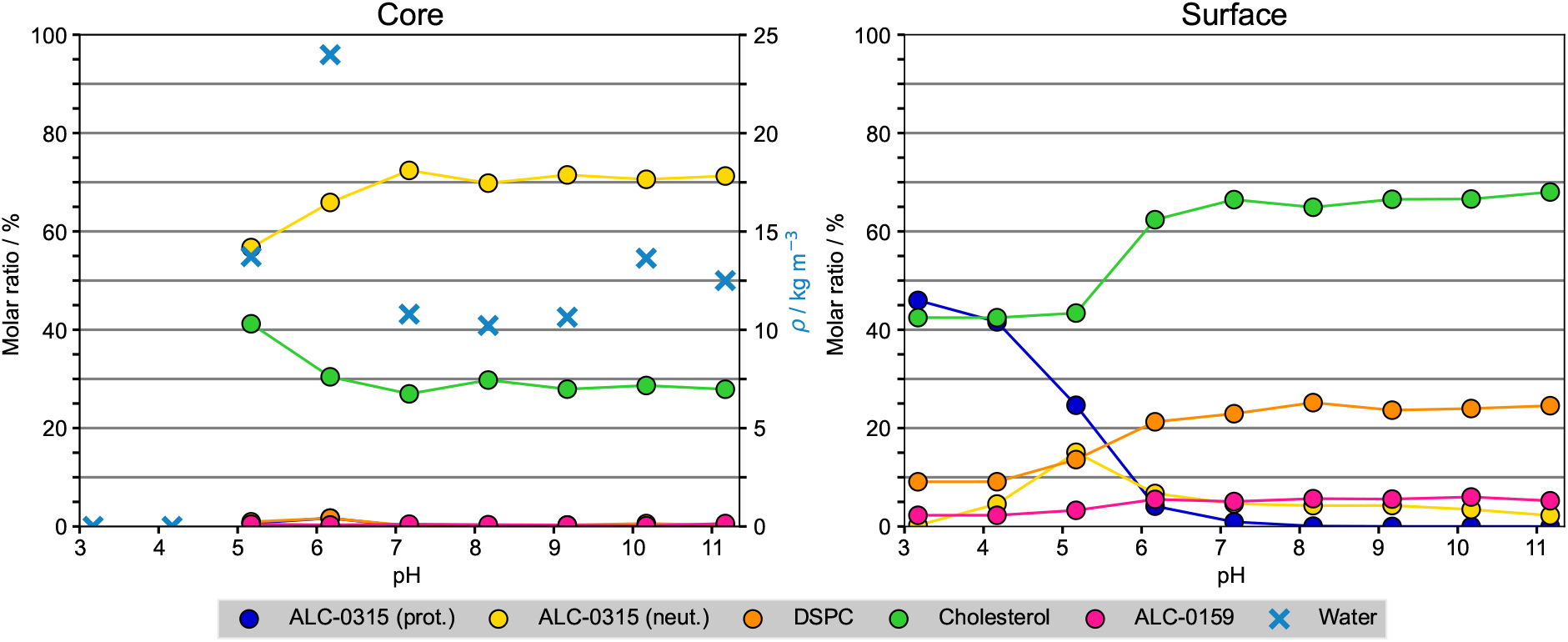
Lipid distribution within LNPs. Molar ratios of lipids within both the core (left panel) and the lipid surface (right panel) of the LNP-mimetic systems as a function of pH are shown. Additionally shown is the mass density of water (*blue*, top panel) within the LNP core. A molecule was assigned to the core if its selected atom was more than 1.8 nm below the LNP surface, defined by the median *z*-position of the phosphorus atoms in DSPC in one leaflet. The following atoms were selected for each molecule type: nitrogen for ALC-0315, phosphorus for DSPC, oxygen for cholesterol, nitrogen for ALC-0159, and oxygen for water.

The CpHMD simulations provide unprecedented insight into the protonation characteristics and pH-driven structural reorganization of the LNP formulation, even under near-physiological pH conditions. Already at pH 5, ≈ 65% of the aminolipids are deprotonated compared to 9% at pH 4, initiating a phase transition characterized by increased membrane thickness as deprotonated aminolipids migrate into the membrane core. This shift reduces the number of protonated aminolipids at the membrane/solvent interface while increasing the density of deprotonated aminolipids within the membrane core (Fig. 1b).

The cholesterol concentration in the monolayers increased significantly from ≈ 42% at pH 3 to ≈ 68% at pH 11 (Fig. 2), close to the cholesterol solubility limit in phospholipid membranes.^44^ Among the helper lipids, DSPC and the PEGylated lipids (ALC-0159) preferentially remain at the membrane surface within the lipid monolayer shell (see Fig. 1b). Occasionally, a few molecules transiently integrate into the membrane core, likely due to favorable interactions with aminolipids during structural rearrangement. The PEGylated lipids may stabilize hydrophobic defects and facilitate the positioning of neutral aminolipids at the membrane surface.^19^

### Interactions between mRNA and lipids

To investigate the impact of mRNA on aminolipid protonation behavior and the spatial distribution of lipids and mRNA, we additionally performed CpHMD simulations at neutral pH of LNP-mimetic systems incorporating mRNA strands.

Figure 3 shows representative snapshots of LNP-mimetic systems containing four mRNA strands of 20 nucleotides each during CpHMD simulations at pH 7, reflecting the *N/P* ratio of the *Comirnaty* formulation. The starting structures for these simulations were derived from microsecond constant-protonation simulations^19^ (details in Methods section).

**Figure 3:**
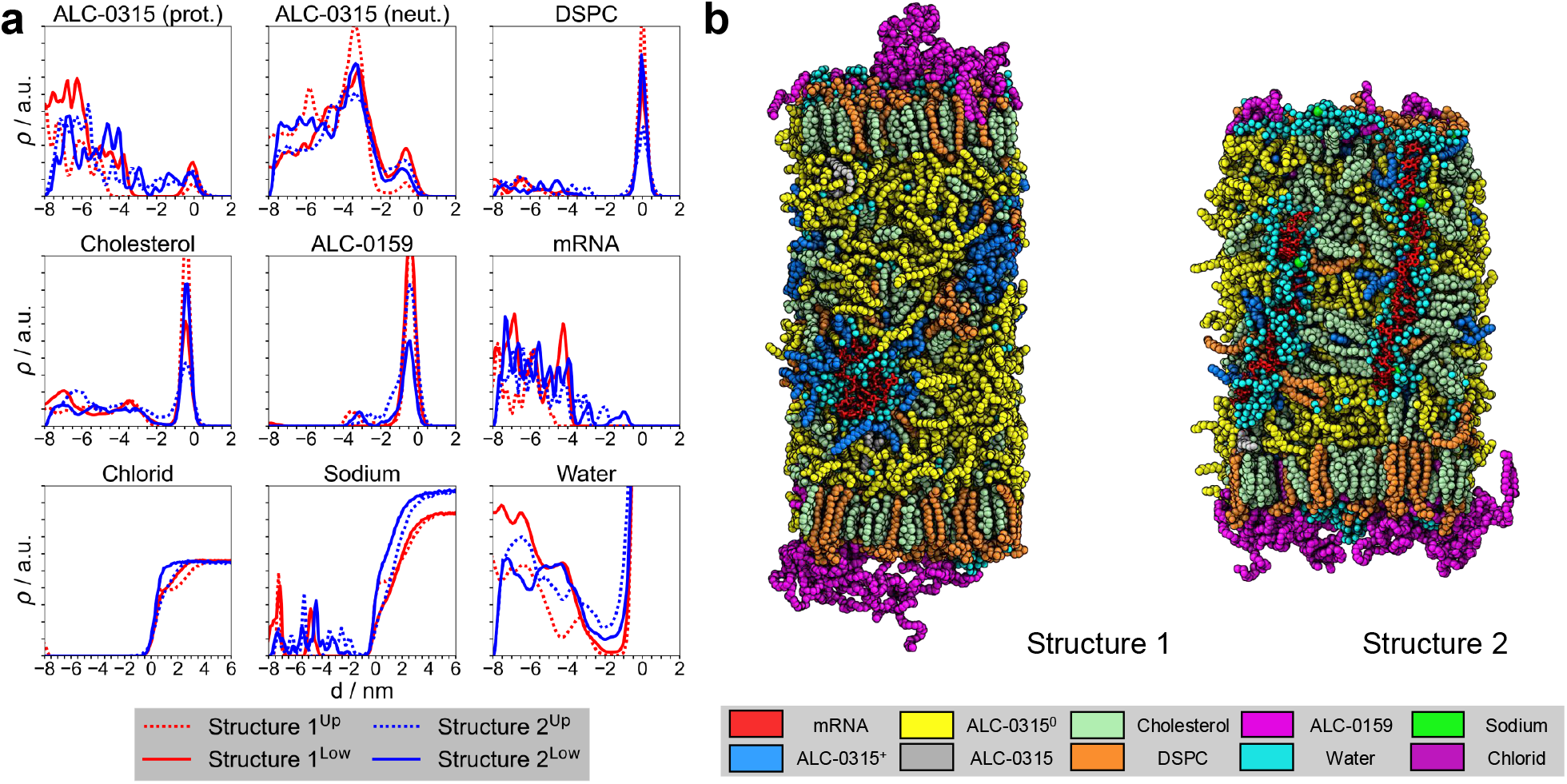
Comirnaty LNP containing mRNA. **a** Average mass densities of all molecule types relative to the membrane surface, defined by the median *z*-position of the phosphorus atoms in DSPC in one leaflet, for two different mRNA configurations (Structure 1: Coil, Structure 2: Elongated) inside the LNP core at pH 7.17. The density curves were calculated using the following atoms for each molecule type: nitrogen for ALC-0315, phosphorus for DSPC, oxygen for cholesterol, nitrogen for ALC-0159, oxygen for water, and phosphorus for mRNA. Density curves represent averages taken after equilibration of *S*^deprot^ values (Supplementary Fig. 9) and were scaled to facilitate comparison between different molecule types. **b** Snapshots of two different mRNA (*red*) configurations (Structure 1: Coil, Structure 2: Elongated) inside the LNP-mimetic system at pH 7.17. The nucleotide strands are embedded within the lipid bulk phase and surrounded by water (*cyan*), protonated ALC-0315 (*blue*), DSPC (*orange*), cholesterol (*palegreen*), sodium ions (*green*), and chloride ions (*purple*). Deprotonated ALC-0315 (*yellow*) is depleted around the nucleotides. The remaining helper lipids, protonated ALC-0315, and ALC-0159 (*magenta*) are located at the surface facing towards the polar solvent. ALC-0315 without a specific protonation state (0.2 ≤ *λ* ≤ 0.8) is shown in *gray*. Images were rendered with PyMOL.^43^

In contrast to mRNA-free systems, CpHMD simulations reveal a substantial presence of protonated aminolipids and water molecules within the LNP core at neutral pH (Fig. 3a). These protonated aminolipids localize near the negatively charged mRNA strands (Supplementary Fig. 13), forming polar cavities within the hydrophobic LNP core. Despite these cavities, enriched with water and cations (Na^+^-ions), the overall LNP structure is maintained, characterized by a hydrophobic core surrounded by lipid monolayers (Fig. 3b).

### *Comirnaty* -formulation lowers the apparent pK_*a*_ of ALC-0315 aminolipid 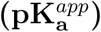

The *apparent* pK_a_ of aminolipids within the *Comirnaty* lipid formulation was derived by fitting the Henderson-Hasselbalch equation (see Eq. 4) to the fraction of deprotonated aminolipids in the LNP-mimetic membrane patch (*S*^deprot^, Fig. 4a).

**Figure 4:**
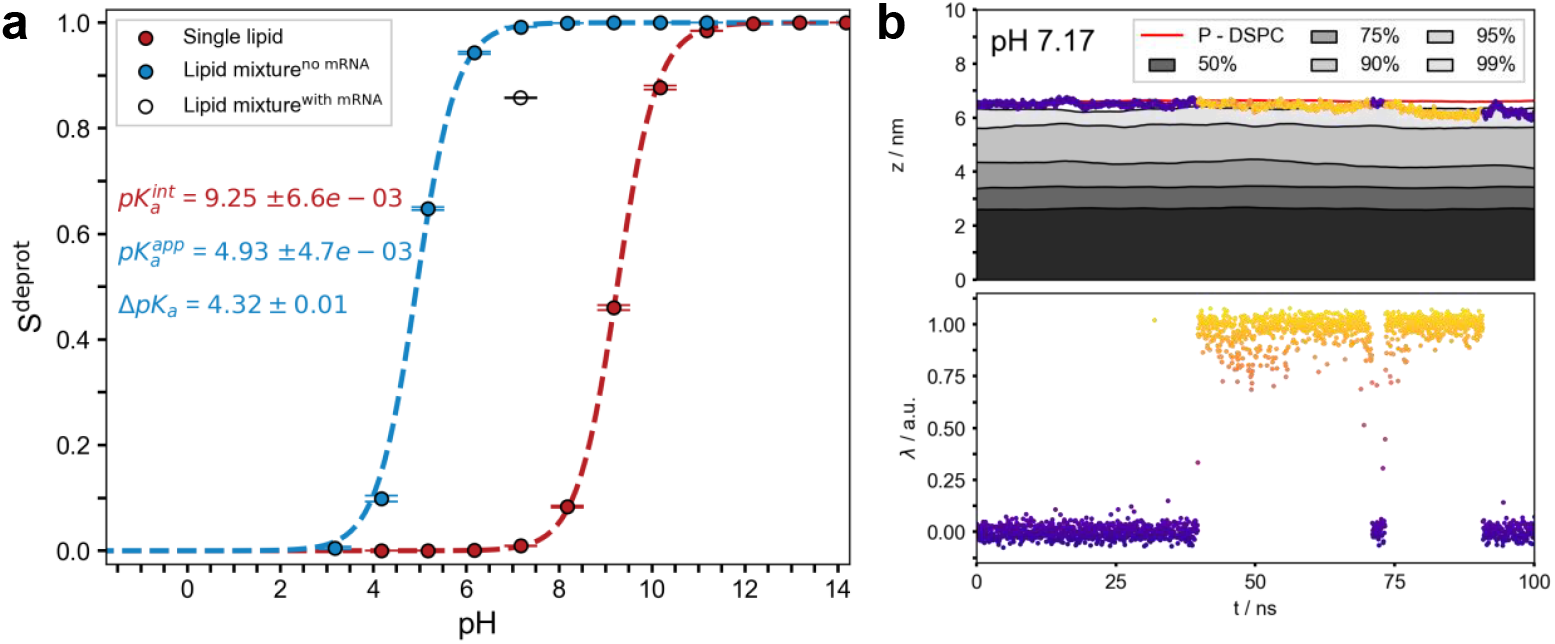
Protonation characteristics of aminolipids. **a** Titration curves of ALC0315 solvated in water (*red*) and within a LNP-mimetic system based on the *Comirnaty* formulation (*blue* without mRNA, and *black* data point with mRNA, Structure 1). Circles indicate the fraction of deprotonated lipids (*S*^deprot^) at each pH. Dashed lines represent fits using the Henderson-Hasselbalch equation to determine the intrinsic and apparent pK_a_ values. Error bars for *S*^deprot^ represent the standard error of the mean, either calculated via block averaging (*blue*)^45^ or from replica simulations (*red*). Averages were calculated after the *S*^deprot^ values reached equilibrium. Errors in the fit parameters were estimated using bootstrapping.^46^ Assuming that *S*^deprot^ is normally distributed at each pH value, new values were sampled from a normal distribution with an expectation value equal to the calculated average and a standard deviation equal to the standard error of the mean. The sampling was repeated 100,000 times; the reported pK_a_ values corresponds to the mean of the bootstrap distribution, and the error bar represents its standard deviation. **b** The upper panel shows the *z*-coordinate of the nitrogen atom in a single aminolipid within the LNP-mimetic over time, colored by its *λ*-coordinate (*blue* ≈0; *yellow* ≈1). Bulk aminolipids are represented by the 50% −99% quantiles of their nitrogen *z*-coordinates, smoothed with a 5 ns running average. The lower panel illustrates the *λ*-coordinate over the same period using the same color scheme. Spatial coordinates are given relative to the membrane center, defined by the median position of the phosphor atoms in DSPC.

The lipid environment induces a significant shift in the apparent 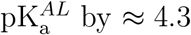 units with respect to the intrinsic pK_a_ of 9.26 to 4.93 ± 0.01 (Fig. 4a), corresponding to a protonation equilibrium shift by ≈ 26 kJ/mol. This apparent 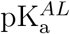 is to be distinguished from the experimentally determined pK_a_ for the LNP of 6.09 obtained using a fluorescence-based assay with 2-(p-toluidino)-6-napthalene sulfonic acid (TNS, see below). ^47^

Fig. 4b illustrates, over a representative time interval and aminolipid, how the *λ*-coordinate, which defines the protonation state of an aminolipid (*λ* = 0 for protonated, *λ* = 1 for deprotonated), is influenced by its position within the LNP-mimetic membrane. When the aminolipid transitions from the solvent-membrane interface to the bulk lipid phase (e.g., between 40 ns and 70 ns in Fig. 4b), it rapidly deprotonates. Conversely, upon returning to the polar membrane interface (e.g., between 70 ns and 75 ns), it promptly protonates. These (de-)protonation events occur on a timescale of just a few nanoseconds, demonstrating the ability of aminolipids to swiftly adapt their protonation state in response to local environmental changes. At the same time, the timescale of protonation adjustments is sufficiently slow to allow for changes in the aminolipid’s membrane localization. However, the combination of comparably fast dynamic de-/protonation events with the slow aminolipid exchange between lipid monolayers and LNP-mimetic core, significantly slows down equilibration of the LNP-mimetic systems in particular for intermediate pH values between 5 and 6 (Supplementary Figures 5–7).

### Position-dependent pK_a_ of aminolipids 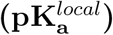

As illustrated in Fig. 4b, the protonation state of aminolipids is strongly modulated by their insertion depth within the LNP-mimetic membrane. Fig. 5a shows that the fraction of deprotonated aminolipids (*S*^deprot^) varies markedly with insertion depth at different pH values. Aminolipids located near the monolayer surface (*d* ≈ 0, defined as the distance from the DSPC phosphorus atoms) display substantially higher protonation levels (i.e., lower *S*^deprot^) compared to those embedded within the LNP core phase (*d <* 0, Fig. 5b), indicating that protonation is energetically favored at the solvent-lipid interface. Remarkably, even at pH ≈ 7, well above the apparent aminolipid pK_a_, a significant fraction (≈ 17 %) of surface aminolipids remains protonated (Fig. 2b).

**Figure 5:**
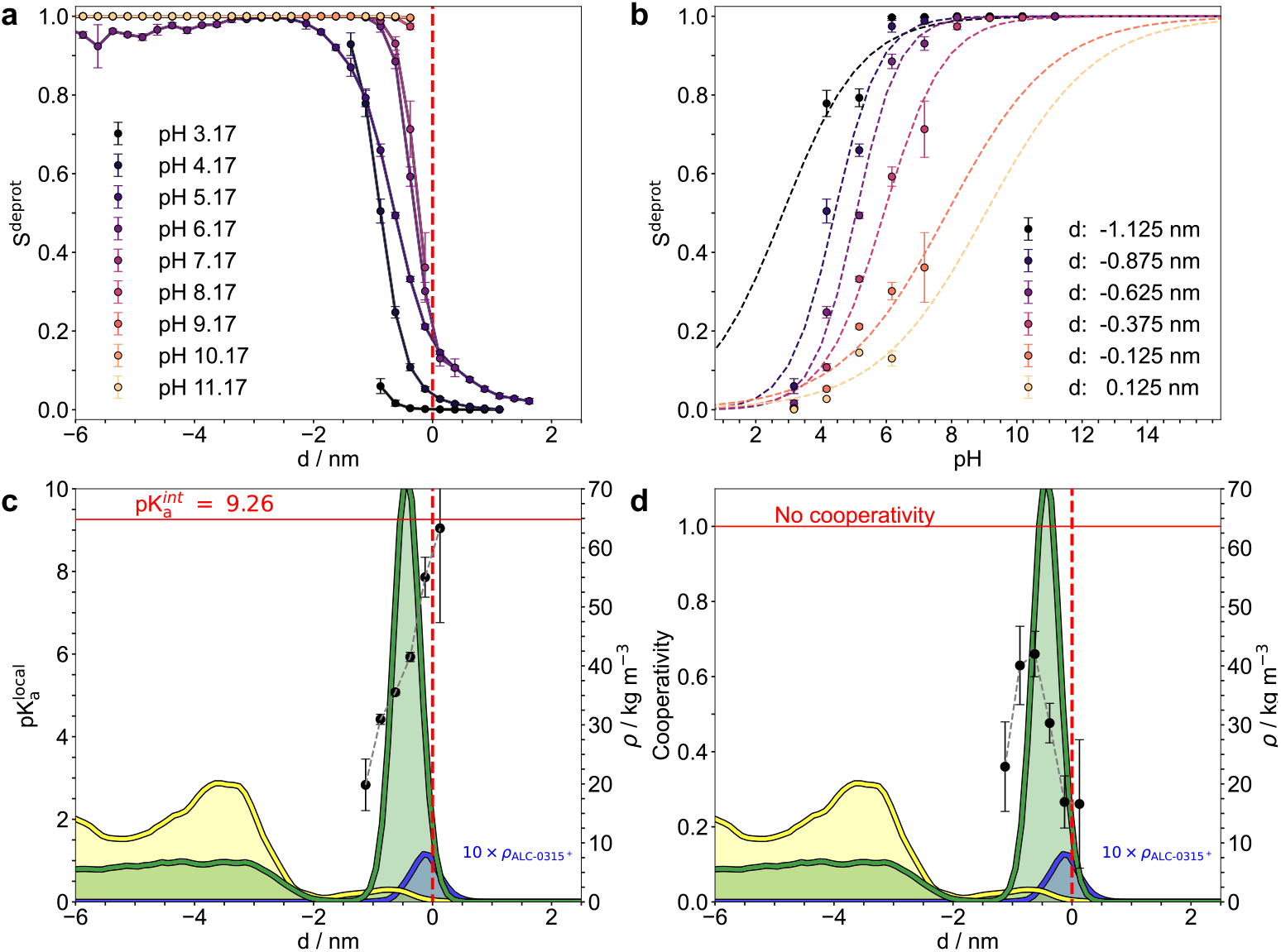
Position-dependent pK_a_ of aminolipids. **a** Fraction of deprotonated ALC-0315 aminolipids (*S*^deprot^) at different pH values, plotted as a function of their distance from the median *z*-position of DSPC phosphorus atoms, averaged over both leaflets (bin width of 0.25 nm). Negative bin centers indicate titratable sites within the membrane, while positive bin centers correspond to sites above the membrane surface. Error bars represent the standard error of the mean calculated via block averaging. ^45^ **b** Local titration curves for ALC-0315 within the membrane patch, showing *S*^deprot^ values averaged over both leaflets. Dashed lines represent fits to the generalized Henderson-Hasselbalch equation (see Eq. 5). Bins with fewer than 3 values or distances below *z* = −1.2 nm or above *z* = 0.2 nm, respectively, were excluded. Error bars respresent the standard error of the mean calculated via block averaging.^45^ **c** Local apparent pK_a_ values 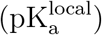 as a function of distance from the membrane surface, and **d** Cooperativity parameter (*n*) from Eq. 5, plotted against the distance to the membrane surface. Error bars represent one standard deviation.^48^ Additionally shown are the lipid densities at pH 7.17 for cholesterol (*green*), deprotonated (*yellow*), and protonated aminolipids (*blue*). Note that the density of the protonated aminolipids was multiplied by a factor of 10 for improved visual representation.

The curves of *S*^deprot^ as a function of insertion depth (Fig. 5a), obtained from CpHMD simulations at different pH levels, were converted into pH-dependent titration curves for distinct insertion depths (Fig. 5b). This transformation enabled the determination of *local apparent* 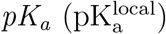 values and *local degrees of cooperativity* by fitting the data with the generalized Henderson-Hasselbalch equation (Eq. 5). The resulting values are plotted as data points in Fig. 5c+d (*bullets*). Incorporating the cooperativity parameter *n* in the analysis assumes that aminolipids behave similarly to polyelectrolytes ^12^ at the sub-nanometer scale.

The apparent 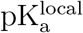 at the monolayer surface of the LNP-mimetic ranges from 6 to 8, approaching the intrinsic pK_a_ of ALC-0315 in pure solvent (*red line*, Fig. 5c). This surface 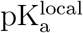 is notably higher than the overall LNP 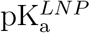 of 6.09. In contrast, at an insertion depth *d* = −1.0 nm, within the cholesterol-depleted region, the local pK_a_ decreases sharply to approximately 4. This pronounced depth dependence is further reflected in the mass density profiles of aminolipids at pH 7 (*yellow* and *blue lines*, Fig. 5c+d). These profiles reveal a preferential localization of protonated ALC-0315 (*blue*) at the monolayer surface, whereas deprotonated aminolipids (*yellow*) are more abundant deeper within the LNP-mimetic membrane patch (*d <* −1.8 nm).

A value *n <* 1 indicates anti-cooperativity in the LNP-mimetic, whereby protonation of one aminolipid decreases the likelihood of protonation in neighboring aminolipids. At the LNP-mimetic surface, the cooperativity reaches its minimum of 0.2 (Fig. 5d), increasing to ≈ 0.6 at an insertion depth of about 1 nm. This depth-dependent variation likely arises from differences in electric shielding (relative permittivity) and the local density of protonated aminolipids within the respective regions. The observed anti-cooperative behavior is consistent with titration curves from *ζ*-potential measurements of various LNP compositions reported by Carrasco *et al*.^12^

### pK_a_ of lipid nanoparticles 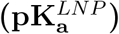

Using the CpHMD approach, we determined the *apparent* pK_a_ of aminolipids within the LNP-mimetic to be 4.93 ± 0.01, representing a decrease by ≈ 4.3 pK_a_ units relative to the intrinsic pK_a_ of the *Comirnaty* aminolipid in solvent. By contrast, the experimentally measured 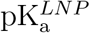 for *Comirnaty* LNPs is 6.09.^49^

How are the intrinsic and apparent aminolipid pK_a_ values related to the overall 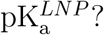 The latter is strongly dependent on the particular experimental method.

In particular, the experimental 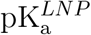 value was obtained using TNS binding assays, which detect fluorescence changes upon TNS partitioning into hydrophobic regions, driven by electrostatic interactions with charged aminolipids. ^50^ Thus, the measured 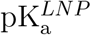 primarily reflects the protonation state of aminolipids accessible to TNS — presumably those located at the LNP surface. This interpretation is consistent with Fig. 2, which shows that charged aminolipids remain enriched at the polar LNP surface up to pH 7.

Considering only aminolipids near the membrane–solvent interface yields a higher calculated pK_a_ of 5.53 ± 0.01 based on a Henderson-Hasselbalch fit to the surface charge density (Supplementary Fig. 14). Including TNS molecules in the simulation system increases this value further to 5.95 ± 0.01 (Fig. 6). This indicates that TNS elevates the measured pK_a_ likely by stabilizing the protonated state of surface aminolipids through electrostatic attraction (via its negatively charged sulfonic acid group).

**Figure 6:**
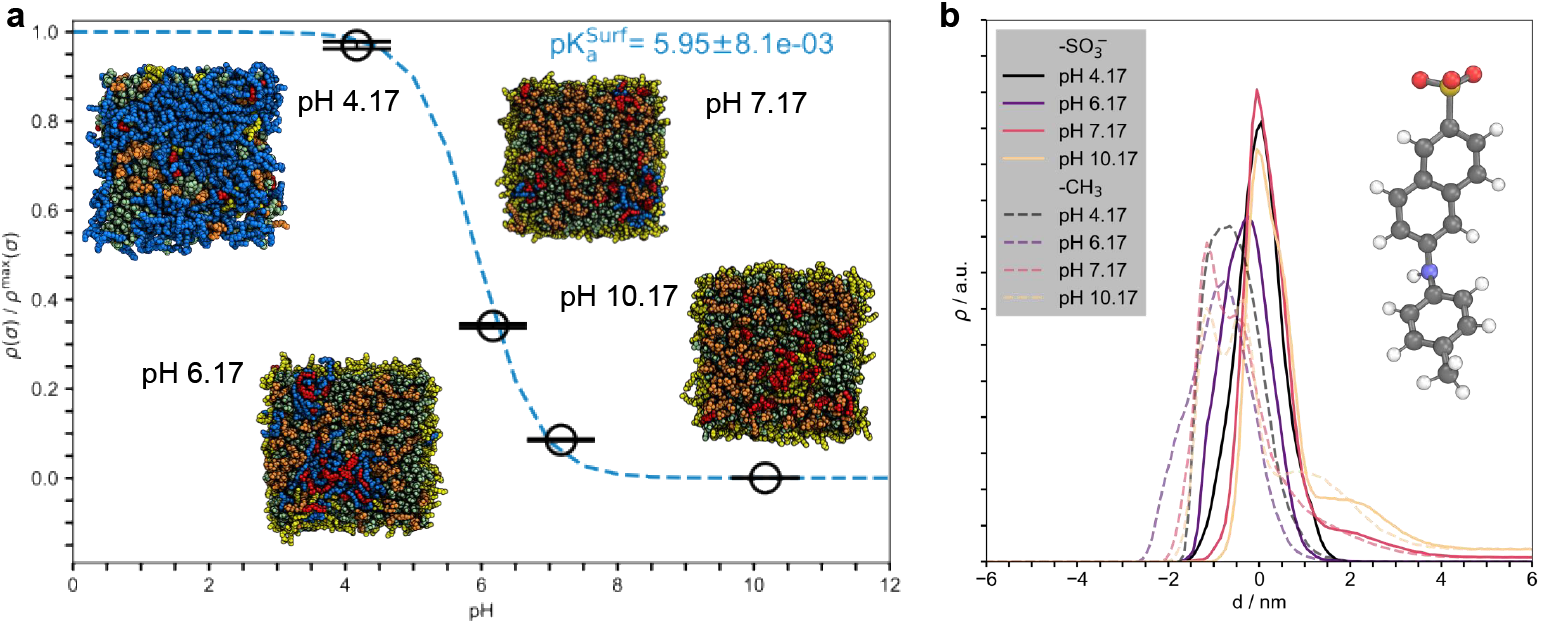
pH-dependent surface charge density of LNP-mimetic systems including the small molecule TNS. **a** The calculation included only protonated ALC-0315 aminolipids within 1.8 nm of, or above, the median position of the DSPC phosphorus atoms in one leaflet. The number of protonations was divided by the respective box area. Charge densities were then averaged over both leaflets in every frame. Error bars represent the standard error of the mean, calculated via block averaging. ^45^ The values were normalized by the maximal charge density, *ρ*^max^(*σ*), derived from system D^‡^ at pH 3. Averages were calculated after the *S*^deprot^ values reached equilibrium. Errors in the fit parameters were estimated using bootstrapping.^46^ Assuming that the scaled surface charge density is normally distributed at each pH value, new values were sampled from a normal distribution with an expectation value equal to the calculated average and a standard deviation equal to the standard error of the mean. The sampling was repeated 100,000 times; the reported pK_a_ values corresponds to the mean of the bootstrap distribution, and the error bar represents its standard deviation. Inset images show the surface of one leaflet of the LNP-mimetic extracted from the last simulation frame: deprotonated ALC-0315 (*yellow*), protonated ALC-0315 (*blue*), DSPC (*orange*), cholesterol (*green*), and TNS (*red*). ALC-0159, and solvent are not shown. **b** Average mass densities of the sulfur and the carbon atom in the head- (“-SO^-^_3_”) and the tail-group (“-CH_3_”) of TNS for each pH level. The densities were scaled to facilitate comparison between the head- and the tail-group. Inset images shows TNS in ball-stick representation (oxygen (*red*), sulfur (*yellow*), carbon (*grey*), nitrogen (*blue*), and hydrogen (*white*). All pictures of atomistic structures were rendered with PyMOL. ^43^

## Discussion

This study provides atomistic insights into the pH-dependent behavior and structural dynamics of LNP-mimetic systems, focusing on aminolipid protonation and its implications for lipid organization and mRNA encapsulation. Using scalable constant-pH MD simulations,^40^ we establish a mechanistic link between the intrinsic pK_a_ of aminolipids in solution and the experimentally observed 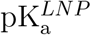 of complete nanoparticles. In contrast to previous MD studies that fixed aminolipid protonation states based on the experimental 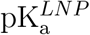 value (≈ 6) for complete lipid nanoparticles^12,49^ — which differs substantially from the intrinsic aminolipid pK_a_ (≈ 9) — our approach allows dynamic protonation switching in response to the local electrostatic environment. The spatial heterogeneity of aminolipid protonation in LNP-mimetic systems is investigated across a physiologically relevant pH range. The absence of a pH gradient between the LNP core and the surrounding solvent justifies the use of CpHMD for these systems.^27,51^

The simulations reveal that increasing pH drives a highly cooperative structural transition from a bilayer-like arrangement at acidic conditions (pH 3 – 4) to an LNP-like architecture with a hydrophobic core and a polar monolayer shell at neutral and basic pH. Assuming a constant area-to-volume ratio, these structures correspond to diameters of 39 – 47 nm for mRNA-free systems and 46 – 54 nm for Comirnaty-like LNPs containing 4,000 nucleotides at neutral pH, in good agreement with experimental values. ^29^ The pH-dependent structural reorganization is accompanied by a redistribution of lipid components: deprotonated aminolipids migrate into the core — composed of ≈ 70% aminolipids and ≈ 30% cholesterol — while cholesterol accumulates within the LNP monolayer shell near its solubility limit (≈ 66% in PC bilayers^44^). Notably, even at neutral pH, a substantial fraction of surface aminolipids remains protonated, potentially facilitating endosomal escape.

From CpHMD simulations, the LNP pK_a_ (including TNS) is determined as pK_a_ = 5.95 ± 0.01, only 0.15 units (0.9 kJ/mol) below the experimental value of 6.09. ^49^ This value must be distinguished from the apparent aminolipid pK_a_: fluorescence-based TNS assays preferentially probe surface protonation, yielding higher protonation levels than predicted for the full LNP, and additionally increased by interactions of the probe with the aminolipids. For instance, at endosomal pH (≈ 5.5^20^), the experimental LNP pK_a_ suggests that 80% of the surface aminolipids are protonated, whereas the CpHMD pK_a_ estimate in absence of TNS (≈ 5.5, Supplementary Fig. 14) yields only 50% protonation. Within the endosomal environment, additional interactions — particularly with the negatively charged endosomal membrane — may further modulate aminolipid protonation. Joint simulation–experimental efforts will therefore be essential to fully resolve the interplay between aminolipid protonation and environmental factors (e.g., the endosomal membrane, proteins).

The apparent pK_a_ of ALC-0315 within the LNP-mimetic was determined to 4.93 ± 0.01, a substantial decrease by ≈ 4.3 units relative to its intrinsic value in solvent, and even more than one unit below the LNP pK_a_. Comparable though smaller shifts (2–3 units) between intrinsic aminolipid pK_a_ and LNP pK_a_ have been reported for other ionizable aminolipids^12^ as well, such as D-Lin-DMA-MC3, highlighting that strong pK_a_ depression is a general feature of LNPs. Simpler computational approaches based on potentials of mean force calculations^52^ likely underestimate this effect, as they neglect pH-driven phase transitions and cooperative interactions.

Local pK_a_ analysis reveals strong dependence on the aminolipid insertion depth within the LNP, with values approaching 7–8 near the LNP-mimetic surface but dropping to *<* 4 in the LNP core. Anti-cooperative effects (*n <* 1) are strong at the interface, likely due to variations in dielectric environment and local charge density. Among experimental methods, in particular *ζ*-potential measurements may capture these cooperative effects. ^12,53^ Indeed, simulations with explicit TNS show that the probe even stabilizes the protonated state of surface aminolipids and thus increases the 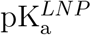 (Figure 6 vs. Supplementary Fig. 14).

Inclusion of mRNA in CpHMD simulations of LNP-mimetic systems further supports the hypothesis that polynucleotides elevate the local pK_a_ of aminolipids, thereby enhancing their protonation within the LNP core^19^ and creating hydrated, ion-containing cavities. This arrangement is consistent with models where mRNA is solvated in aqueous cylindrical channels^23^ rather than fully embedded in an apolar lipid phase.^27^ Notably, the 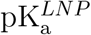 analyzed based on the LNP surface charge is hardly affected by the presence of mRNA (Supplementary Fig. 14, data point marked by *red* circle).

Together, these findings demonstrate that aminolipid protonation equilibria are governed by the local lipid environment, cooperative effects, and cargo-induced effects. By linking molecular-level protonation dynamics to macroscopic observables such as LNP pK_a_ and size, our work provides a mechanistic basis for understanding LNP stability and functionality. These insights into aminolipid localization, protonation, and cargo interactions establish a foundation for the rational design of next-generation LNP formulations with tunable pH responses, enhanced stability, and improved therapeutic performance.

## Methods

### Force field parameters for ALC-0315 (prot/neut) and ALC-0159

Force field parameters for ALC-0315 and ALC-0159 were retrieved from the CHARMM General Force Field (CGenFF v4.6)^54^ with the CGenFF program (v3.0).^55–57^ The partial charges of the atoms within the acyl chains, PEG monomers, ester bonds, and hydroxyl groups were manually set to default values of CGenFF^54^ and the CHARMM force field for lipids.^58^ For the titratable group in the aminolipid, the partial charges used for both the protonated (charge: +1.0) and the deprotonated (charge: 0.0) state were assigned by the CGenFF program.^55^ In the protonated state, a remaining charge of 0.003 e was added to the proton resulting in a total charge of *q*^*H*^ = 0.323 e. The partial charges in the linker region between the hydrophobic tails and the polymer of the PEG-ylated lipid were assigned based on the parameters for dimethylacetamide, which was already included in CGenFF (resname: DMA).^54^ Atom types or bonded potentials assigned by the CGenFF program^55–57^ were not modified. Cholesterol and DSPC parameters were taken from the CHARMM36 force field for lipids.^58–60^ The July 2022 version of the CHARMM36 force field port for Gromacs was used for all simulations.

### Constant-pH parameters for ALC-0315

The influence of the pH on a realistic LNP lipid formulation was studied using the recently published constant-pH code implemented in Gromacs by Aho *et al*.^40^ The algorithm uses *λ*-dynamics to interpolate between charges corresponding either to the protonated (*λ* = 0) or the deprotonated (*λ* = 1) state of each titratable group. The *λ*-dependent electrostatic potential Φ(**R**_*i*_, *λ*) on atom *i* is then defined as:

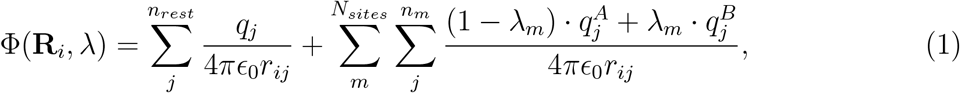

with *n*_*rest*_ as number of atoms not part of any titratable site; *N*_*sites*_ as number of titratable sites; and *n*_*m*_ as number of atoms part of a titratable site each with a partial charge of *q*_*j*_. By interpolating partial charges instead of potential energy functions, the computational overhead is significantly reduced, allowing the simulation of hundreds of titratable sites within the same system. ^40^ Here, the LNP-mimetic systems typically contain 488 titratable aminolipids (see Table 1). Additionally, three *λ*-dependent potential energy functions are added to the total Hamiltonian of the system: the correction potential (*V* ^*MM*^ (*λ*)), the biasing potential (*V* ^*bias*^(*λ*)), and the pH-dependent potential (*V* ^*pH*^ (*λ*)).^40^ The terms *V* ^*bias*^(*λ*) and *V* ^*pH*^ (*λ*) are analytical functions that are described in more detail in Aho *et al*.^40^ If not otherwise specified, the simulations in this paper applied a biasing potential *V* ^*bias*^(*λ*) with a barrier height of 7.5 kJ mol^-1^, corresponding to the default value set by pHbuilder.^61^ The correction potential *V* ^*MM*^ (*λ*) is defined as the negative of the deprotonation free energy,

**Table 1:**
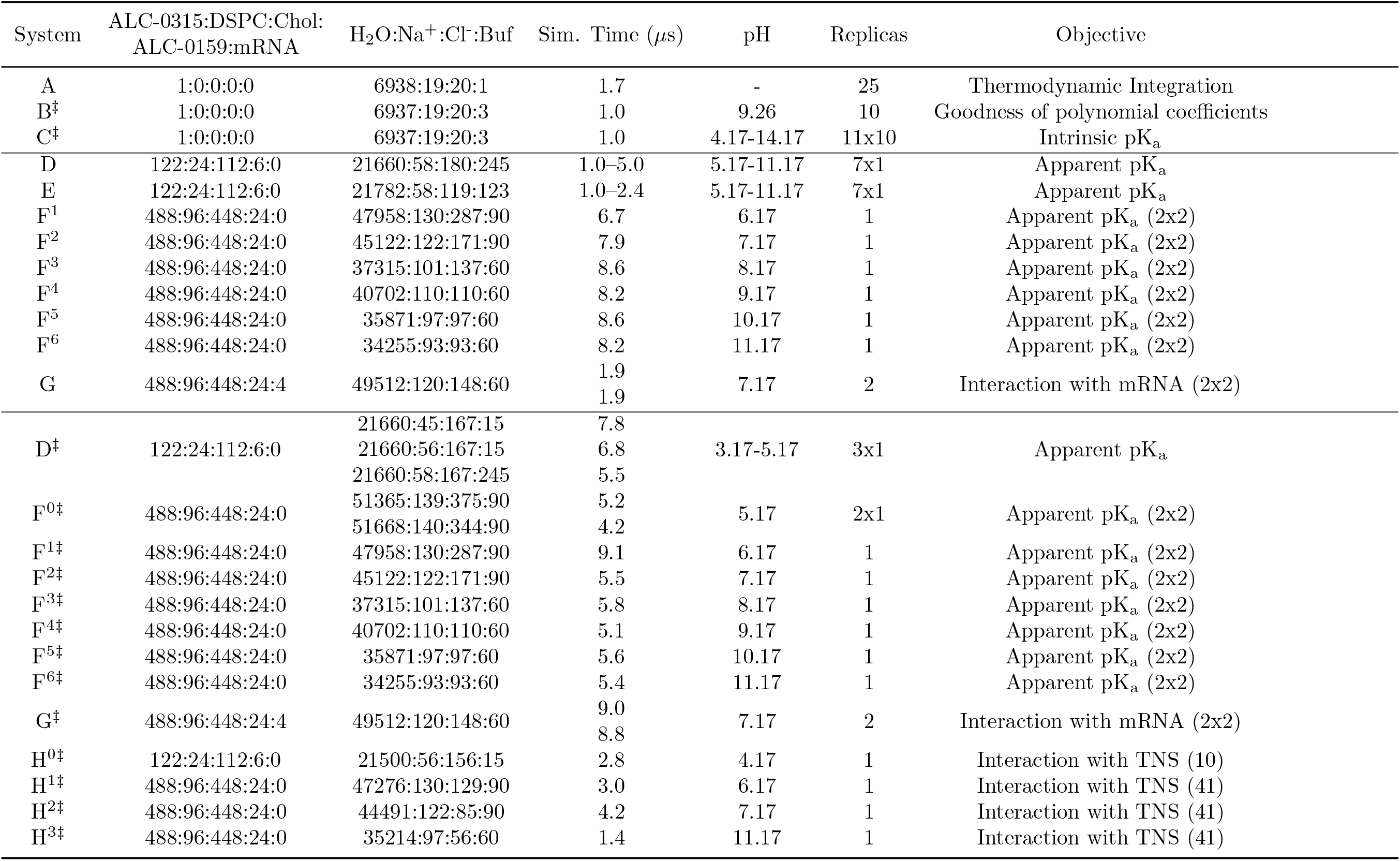
Systems studied with all-atom molecular dynamics simulations using the constant-pH method. The number of solvent molecules represents always the final topology. (^†^ALC-0315^Swap, ‡^ALC-0315^Corrected^)

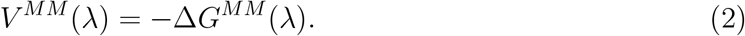

The implementation of Aho *et al*.^40^ uses a polynom to represent *V* ^*MM*^ (*λ*). We employed the workflow described by Buslaev *et al*.^62^ and Jansen *et al*.^61^ to derive the *a priori* unknown polynomial coefficients via thermodynamic integration. The modified Gromacs version^40^ and the pHbuilder tool (v1.2.4/5 & v1.3)^61^ were used to generate all relevant structural and initial topology files for the parameterization. For the thermodynamic integration (system A in Table 1), a single aminolipid was initially placed in a 6×6×6 nm^3^ box solvated with water, one counter ion, and an additional salt concentration of 150 mM Na^+^Cl^-^. During the simulation, position restraints were applied to avoid strong interactions between the aminolipid and the buffer particle. The potential energy of the initial structure was minimized using the steepest descent algorithm until a tolerance of 1,000 kJ mol^-1^ nm^-1^ or a maximum of 5,000 steps was reached (maximum step size: 0.01 nm). Following energy minimization, the system was equilibrated in the NVT ensemble for 100 ps, and subsequently in the NPT ensemble for 900 ps, using the leap-frog integrator with a time step of 0.001 ps and 0.002 ps, respectively. The equilibrated structure then served as the initial configuration for the different *λ* windows. No initial velocities were generated at the start of the simulation.

In general, temperature and pressure were maintained in all simulations using the velocity-rescale algorithm^63^ (T^ref^=310 K and *τ*_*T*_ = 0.5 ps) and the C-rescale algorithm^64^ (p^ref^=1 bar, *τ*_*P*_ = 5.0 ps, and a compressibility of 4.5 × 10^−5^ bar^-1^), respectively. Both quantities were coupled with a frequency of 20 steps. According to the recommendations of Kim *et al*.^65^ the automated buffer setting was overridden (Verlet-buffer-tolerance = -1) and the outer cut-off distance for the short-range neighbor list (rlist) was set to 1.35 nm with a frequency of 20 steps to update the neighbor list (nstlist). Forces from van der Waals interactions were smoothly switched to 0 between 1.0 nm and 1.2 nm (inner cut-off distance). Electrostatic interactions were handled using the fast smooth Particle-Mesh Ewald (PME) method^66^ with a real-space cut-off distance of 1.2 nm and a Fourier spacing of 0.14 nm. Bonds including hydrogen atoms, were constrained and handled with the LINCS algorithm. ^67^

For system A (see Tab. 1), the average minimum distance between all atoms in the aminolipid and the buffer particle was 3.54 nm, with the distances ranging from a minimum of 2.48 nm to a maximum of 4.34 nm. Contributions from the three constant-pH specific potentials *V* ^*MM*^ (*λ*), *V* ^*bias*^(*λ*), and *V* ^*pH*^ (*λ*), are not considered during the thermodynamic integration simulations. Averages of 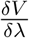 were sampled for *λ* values between −0.10 and 1.10 with an equidistant spacing of 0.05 *λ*, and a temporal output frequency of 1 ps. Estimates of 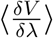 were obtained from 100 ns to 1700 ns of each *λ* window simulation. Convergence of the estimate was checked by monitoring the value of 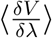 over time (see Supplementary Fig. 1a). Polynomial coefficients were obtained by a least-squares polynomial fit using *polyfit* from NumPy v2.0.0^68^ (see Supplementary Fig. 1b). For the aminolipid, a one-dimensional polynom of order 9 was used to fit 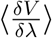.

The quality of the fitted parameters was assessed by running 10 independent simulations (system B, Tab. 1) without position restraints and without contributions from *V* ^*bias*^(*λ*) and *V* ^*pH*^ (*λ*) (i.e., pH = pK_a_). The initial structure of a single aminolipid in solvent was first minimized and then equilibrated for each replica simulation, using the three-step protocol described above. Initial velocities were not generated for the replica simulations. The distribution of *λ* should then obey 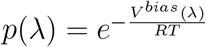 under these conditions if a proper estimate of the correction potential is provided^40,61,62^ (see Supplementary Fig. 2).

### Modeling the intrinsic pK_a_ of the aminolipid ALC-0315

The parameterization of ALC-0315 was validated using CpHMD simulations of a single aminolipid solvated in water at physiological salt conditions (150 mM Na^+^Cl^−^) across a pH range of 4 to 14 (Tab. 1, system C). In these simulations, all correction potentials (*V* ^*MM*^ (*λ*), *V* ^*bias*^(*λ*), and *V* ^*pH*^ (*λ*)) were applied, with the pH gradually increasing. The structure was first minimized and then equilibrated again using the three-step protocol described above. The simulations comprised 11 distinct pH conditions (same initial configuration), each with 10 replicas, for a total simulation time of 1.0 *µ*s per replica. Systems were set up using the *create titration*.*py* script provided by pHbuilder.^61^ Unless otherwise specified, the buffer parameters from Buslaev *et al*.^62^ were used.

Any significant deviation of the numerically derived pK_a_ from the pre-defined reference value would suggest either insufficient sampling during titration or an inadequate estimation of the protonation free energy *V* ^*MM*^ (*λ*) via thermodynamic integration. The intrinsic pK_a_ of 9.25 ± 0.01 was determined by fitting the Henderson-Hasselbalch equation (Eq. 4) to the mean aminolipid deprotonation fraction (*S*^deprot^), as shown in Fig. 4a. This value excellently matches the reference pK_a_ of 9.26,^52^ indicating both sufficient sampling and accurate parameterization of the protonation behavior of ALC-0315.

### Force field parameters for 2-(p-toluidino)-6-naphtalene sulfonic acid (TNS)

Force field parameters for 2-(p-toluidino)-6-naphtalene sulfonic acid (TNS) were retrieved from the CHARMM General Force Field (CGenFF v4.6)^54^ with the CGenFF program (v4.0).^55–57^ Penalties for both non-bonded and bonded parameters were equal to 0.0, so the obtained parameters were used in the subsequent simulations without further modifications. In all subsequent simulations, TNS was assigned a constant net charge of –1.

### Multi-component systems

#### Lipid-Only Systems

The CpHMD simulations of the lipid composition of *Comirnaty* LNPs were started from an equilibrated snapshot at low pH previously obtained in classical MD simulations using a fixed protonation for the aminolipids.^19^ The snapshot was taken after 4 *µ*s of an all-atom simulation. The molar lipid ratio for ALC-0315:DSPC:cholesterol:ALC-0159 (PEGylated lipid) is 46.3:9.4:42.7:1.6, corresponding to 122 aminolipids, 24 DSPC molecules, 112 cholesterol molecules, and 6 ALC-0159 (PEGylated lipids) (systems D and E, Table 1).

Prior to starting the CpHMD simulations, an energy minimization was performed, with a maximum step of 0.0001 nm, an energy tolerance of 1,000 kJ mol^-1^ nm^-1^, a maximum of 5,000 steps, and fixed protonation states. CpHMD simulations were performed in the pH range between 5.17 … 11.17 with a step size of 1 pH unit. To probe the influence of different buffer particle concentrations and different initial protonation states of the lipids, we repeated the titration for two different initial states (system D, E, Tab. 1). In system D (basis system for subsequent simulations), 245 buffer atoms were added with all lipids starting from a protonated state. The number of buffer atoms equals 2·*N* +1 with *N* representing the number of titratable residues in the system. In system E, 123 buffer atoms were added with only half of the lipids initially in their protonated state. Simulations ran for a minimum of 1 *µ*s. No significant differences were observed for the length of the box vectors (Supplementary Fig. 3) and the number of (de)protonated lipids (Supplementary Fig. 4) between systems with differing buffer particle size. However, at pH 6.17, which is close to the apparent pK_a_, the membrane surface area decreased more rapidly in system D (higher buffer concentration). The mass density profiles shown in Supplementary Fig. 12 indicate that the buffer particles avoid the hydrophobic interior of the membranes.

Enlarged systems were setup by quadruplicating snapshots of system D (see Tab. 1) in the *x*-*y* plane (systems F^1-6^, Tab. 1). A part of the bulk water was removed to prevent large computational overheads. Remaining ion numbers were adapted to keep the ion concentration comparable between systems. The aminolipid protonation states were kept for the enlarged membranes. Protonation was assigned if the *λ* coordinate was below a value of 0.2.

The amount of buffer atoms was set to 60 particles per system based on the charge fluctuations at pH 7.17. An energy minimization (steepest descent) with a maximum step size of 0.01 nm, an energy tolerance of 1,000 kJ mol^-1^ nm^-1^, and a maximum of 30 steps was performed before starting the simulations of the quadruplicated systems. The systems were re-initialized (protonation states, number of buffer particles, number of ions) if the *λ*-coordinate exceeded the threshold of 1.15. Tab. 1 shows the final molecule numbers for systems F^1-6^.

After extended simulation time, the constant-pH parameters of the aminolipid ALC-0315 were re-evaluated and re-parameterized to ensure correct charge assignment, which was confirmed by single-point energy calculations. Simulations of the complex lipid mixture, with and without mRNA (see below), were restarted from structures and protonation states obtained from the previous trajectories (see Tab. 1). Unless stated otherwise, all results presented are based on simulations performed with the validated parameter set for ALC-0315 (systems marked by ^‡^; systems D^‡^ at pH 3.17-5.17 started from equilibrated basis system D, systems F^0-6‡^ at pH 5.17-11.17 from equilibrated quadruplicated systems F).

#### ipid/mRNA Systems

To investigate the impact of mRNA on the protonation state of ALC-0315 within the LNP core, two equilibrated snapshots of mRNA-lipid simulation systems, previously published by Trollmann and Böckmann,^19^ were used as initial structures. These structures contained four strands of negatively charged poly-nucleotides encapsulated within the lipid bulk phase, which comprised a mixture of protonated and deprotonated aminolipids. Notably, the mRNA strands differ in structure between the two systems as a result of the random self-assembly process used for system generation.^19^ The constant protonation states in these snapshots were taken as initial protonation states of the aminolipids in the CpHMD simulations to preserve the pre-equilibrated interaction network between mRNA and lipids as much as possible. After adding the buffer particles, an energy minimization with a maximum step size of 0.01 nm, an energy tolerance of 1,000 kJ mol^-1^ nm^-1^, and a maximum of 30 steps was performed before starting the simulation. Both systems (system G, Tab. 1) were simulated with a pH of 7.17. Parameters for the (modified) mRNA nucleotides were taken from the CHARMM36 force field.^69,70^

### Lipid/TNS System

Snapshots from systems F^0,2,3,6‡^ (see Tab. 1) at 4.6 *µ*s, 4.5 *µ*s, 5.6 *µ*s, and 4.5 *µ*s were extracted as the initial structures for TNS addition (systems H^0–3‡^). TNS molecules were added to achieve a TNS:aminolipid ratio of 1:12, consistent with experimental assays.^49,50^ The TNS molecules were placed in random orientations on a square grid in the xy-plane with 1 nm spacing between their centers. Their initial z-coordinates were randomly sampled near the maximum length of the z-axis to maximize the distance from the LNP-mimetic membrane surface. Aminolipid protonation states were assigned as described previously (protonated if *λ <* 0.2). This was followed by energy minimization using the steepest-descent algorithm, with a maximum step size of 0.01 nm, an energy tolerance of 1,000 kJ mol^-1^ nm^-1^, and up to 30 steps before beginning production runs. Supplementary Fig. 10-–11 show the box length in the x-direction and the values of *S*^deprot^ over the course of the production simulations.

### Analysis of the constant-pH trajectories

The analysis of the titratable groups follows the recommendations outlined by Aho *et al*.^40^

The fraction of deprotonated aminolipids (*S*^deprot^) was computed by

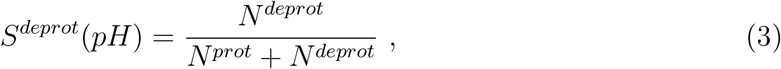

where *N* ^prot^ and *N* ^deprot^ denote, for a single-molecule simulation, the number of frames in which a titratable site is protonated (*λ <* 0.2) or deprotonated (*λ >* 0.8). For systems containing multiple titratable sites, they denote the number of sites that are protonated or deprotonated, respectively.

The intrinsic or apparent pK_a_ was determined by fitting the Henderson–Hasselbalch equation to the calculated values of *S*^deprot^:

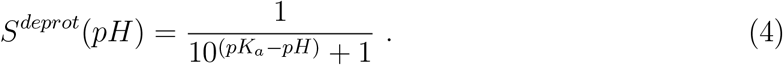

For the analysis of the position-dependent pK_a_ (see Fig.5), the values of *S*^deprot^ were fitted to the generalized Henderson–Hasselbalch equation, as originally applied to titration curves of polymeric acids:^71^

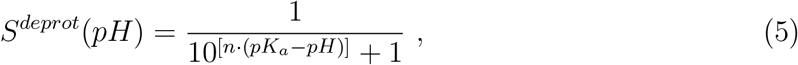

the coefficient *n* was interpreted in this work as a measure of the cooperativity for the protonation within a system of several aminolipids. A value *n <* 1 refers to negative cooperativity, or anti-cooperativity, while *n >* 1 indicates positive cooperativity between the titratable sites. The same function was used by Carrasco *et al*.^12^ to fit titration curves of LNPs obtained from *ζ*-potentials.

### Assessment of simulation convergence and error estimation

Convergence was assessed by monitoring the time evolution of *S*^deprot^ for each simulation. The drift in this observable was quantified by performing a linear regression on segments of the trajectory and computing the difference between the first and last values of the regression model. To obtain a series of drift estimates, the starting point of the regression was incrementally increased (i.e., by steps of 5 ns)—effectively removing the initial part of the trajectory. Each drift value was normalized by the empirical standard deviation of the observable within that block. Equilibration was defined as the point at which this ratio first fell below 1.0. Mean values and error bars were then calculated over the resulting equilibrated range. For time series exhibiting strong drift and large standard deviations, where the ratio also fell below 1.0, the equilibrated range was manually adjusted. Error bars represent the standard error of the mean using block averaging, with fits performed using either the single or double exponential function proposed by Hess. ^45^ Mean values for system F^0‡^(see Tab. 1) were averaged over the two independent simulations, and the standard error of the mean was calculated using standard error propagation rules.

## Acknowledgement

The authors gratefully acknowledge the scientific support and HPC resources provided by the Erlangen National High Performance Computing Center (NHR@FAU) of the Friedrich-Alexander-Universität Erlangen-Nürnberg (FAU). The hardware is funded by the German Research Foundation (DFG).

## Additional information

The supplementary material includes

- Supplementary Figures 1 – 14

## TOC Graphic

## Notes

### Competing Interest Statement

The authors have declared no competing interest.

### Summary of Updates

- The manuscript was streamlined. - We have included substantial new simulation data that expand both the scope and robustness of the study. Specifically, we extended the simulation timescales to better capture slow structural transitions and ensure convergence of the results. In addition, we performed new simulations that explicitly include the fluorescent probe TNS, which is commonly used in experimental assays to determine apparent LNP pKa values.

