## Supplementary for "Decoding pH-Driven Phase Transition of Lipid Nanoparticles"

August 22, 2025

#### The PDF file includes

- Supplementary Figures 1 – 14,
- Supplementary References

### Supplementary Figures

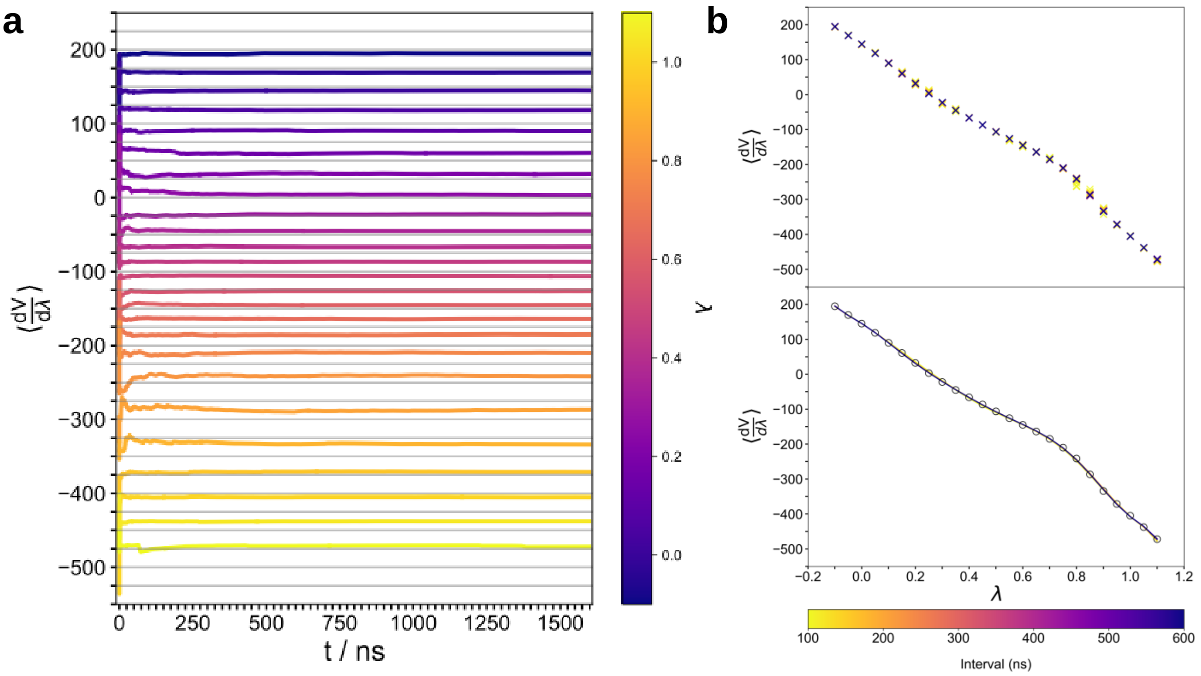


**Figure 1: a** Cumulative estimation of $\langle \frac{dV}{d\lambda} \rangle$ as a function of increasing simulation length. **b** Cumulative estimation of $\langle \frac{dV}{d\lambda} \rangle$ as a function of $\lambda$ (upper panel), and the development of the 9th-order polynomial fit for $\langle \frac{dV}{d\lambda} \rangle$ under an increasing simulation length (bottom panel). The black circles in the bottom panel represent the final estimate for $\langle \frac{dV}{d\lambda} \rangle$ using the whole equilibrated range of the trajectory from 100 ns to 1700 ns.

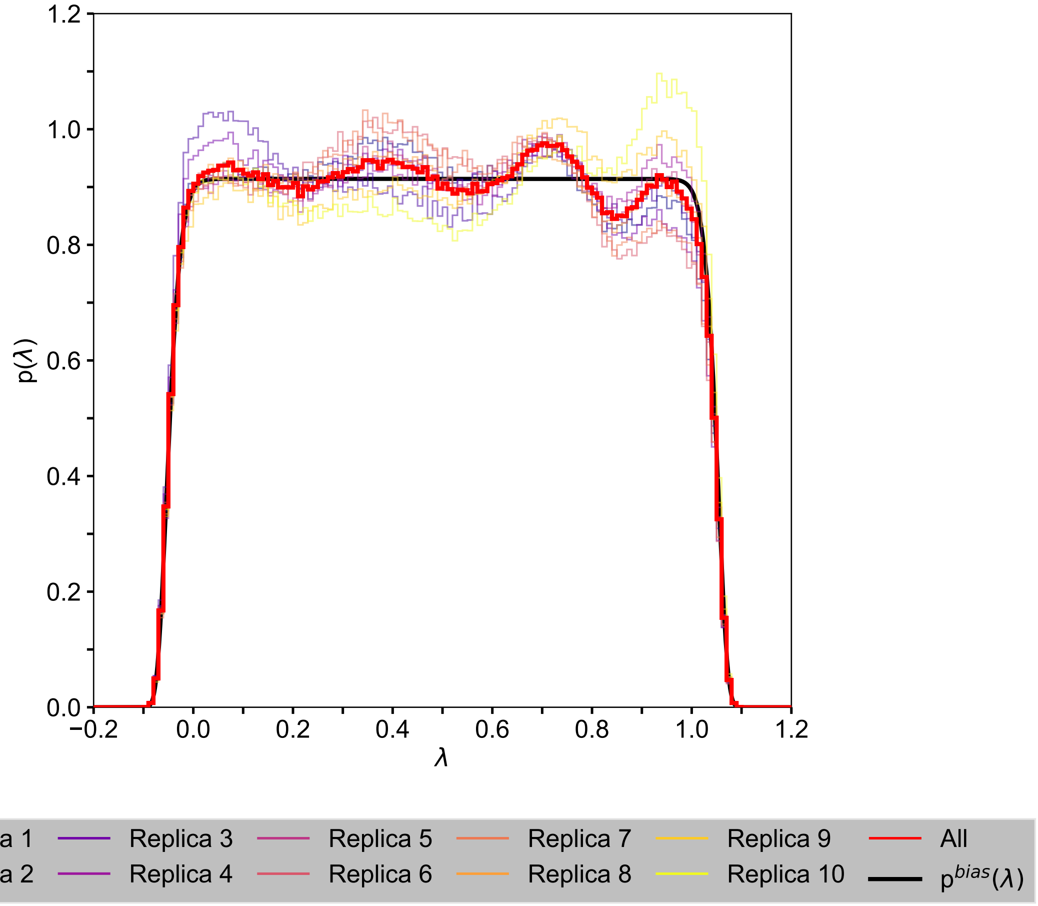


**Figure 2:** Distribution of the $\lambda$-coordinate for the aminolipid obtained from ten independent simulations applying only the correction potential $V^{MM}(\lambda)$ for the protonation free energy and the $V^{\text{bias}}(\lambda)$ (barrier height set to 0.0 kJ $\text{mol}^{-1}$). The expected probability density function, $p^{\text{bias}}(\lambda)$, shown for reference, is derived from the applied bias potential, $V^{\text{bias}}(\lambda)$.

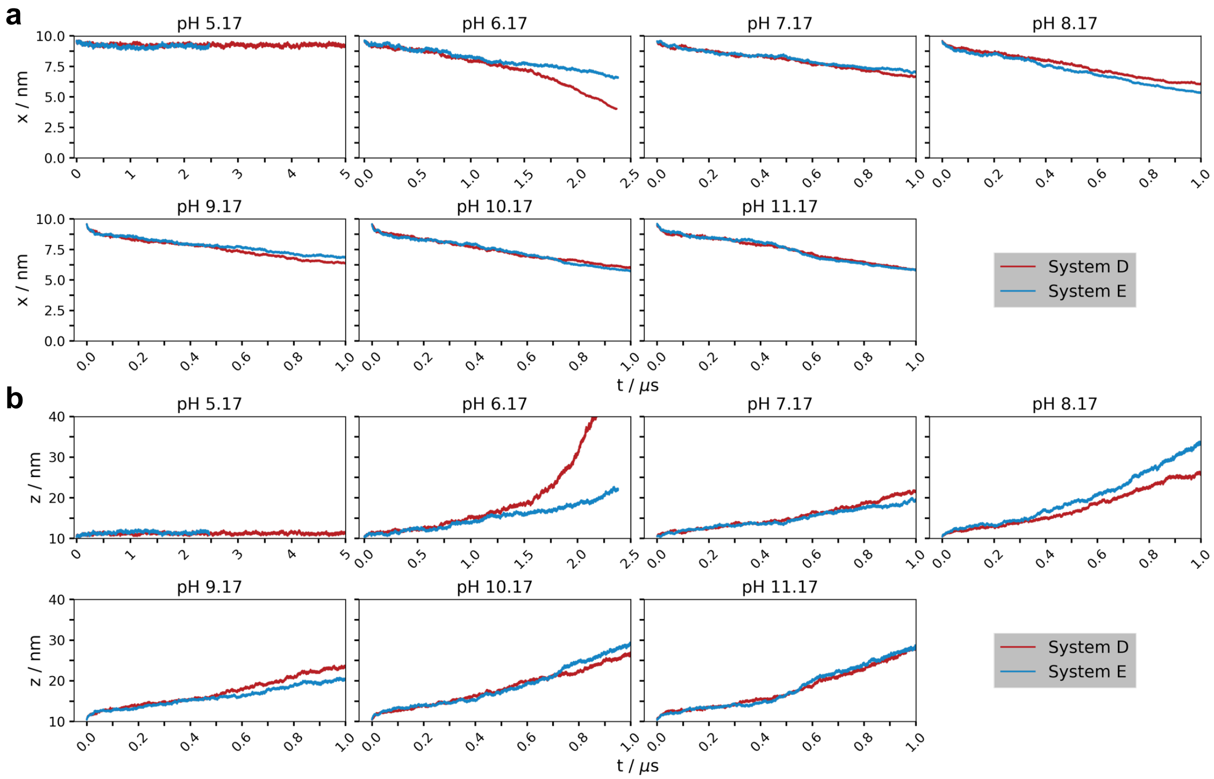


**Figure 3:** Length of the box vectors in **a** the x-direction and **b** the z-direction over time for lipid mixture simulations using either $2 \cdot N + 1$ buffer particles for $N$ titratable sites (System D, Tab. 1) or half that number (System E, Tab. 1).

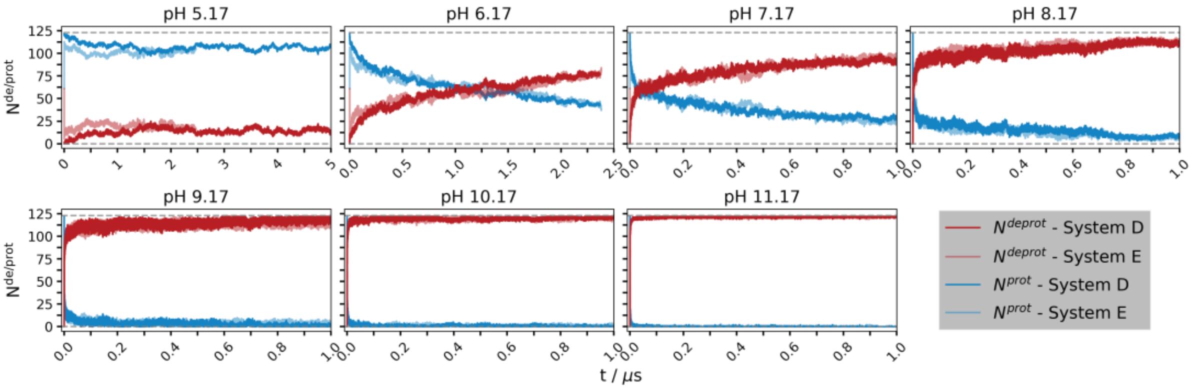


**Figure 4:** Number of (de)protonated residues in lipid mixture simulations using either $2 \cdot N + 1$ buffer particles for $N$ titratable sites (System D, Tab. 1) or half that number (System E, Tab. 1). The number of (de)protonated residues was calculated every 1 ps based on the $\lambda$-coordinate for each aminolipid ($\lambda < 0.2$, protonated; $\lambda > 0.8$, deprotonated).

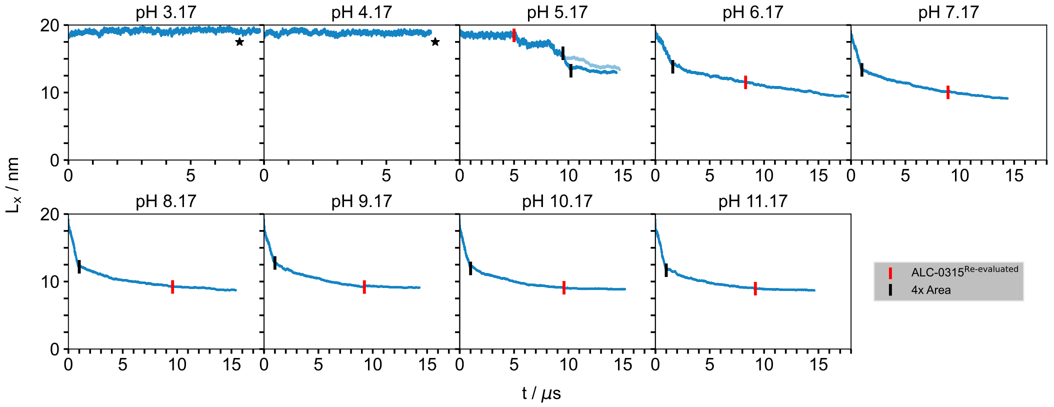


**Figure 5:** Length of the box vector in the x-direction over time for simulations of systems $D_{\text{pH 5.17}}$, $F^{1-6}$, $D^{\ddagger}$, and $F^{0-6\ddagger}$(see Tab. 1). Black vertical lines indicate the quadrupling of the membrane surface, performed to prevent periodic boundary artifacts caused by an undersized surface area, while red vertical lines indicate the restart of the simulations with corrected polynomial coefficients (ALC-0315$^{\text{Re-evaluated}}$). Box lengths prior to the black vertical bar, or annotated with a star ($\star$), were doubled to produce a continuous curve in the plot.

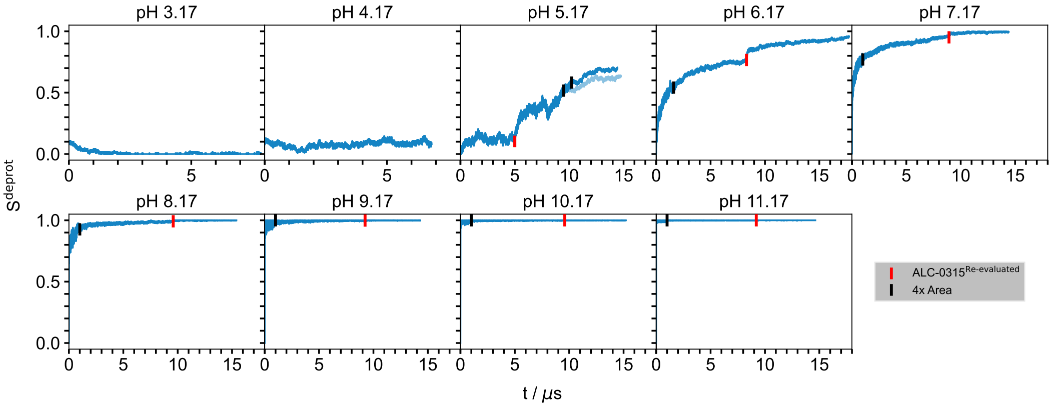


**Figure 6:** Fraction of deprotonation ($S^{\text{deprot}}$) (see Eq. 3) over time is shown for simulations of $D_{\text{pH 5.17}}$, $F^{1-6}$, $D^{\ddagger}$, and $F^{0-6\ddagger}$ (see Tab. 1). The number of (de)protonated aminolipids was obtained via the $\lambda$-coordinate of each aminolipid ($\lambda < 0.2$, protonated; $\lambda > 0.8$, deprotonated) every 1 ps. Black vertical lines indicate the quadrupling of the membrane surface, performed to prevent periodic boundary artifacts caused by an undersized surface area, while red vertical lines indicate the restart of the simulations with corrected polynomial coefficients (ALC-0315$^{\text{Re-evaluated}}$).

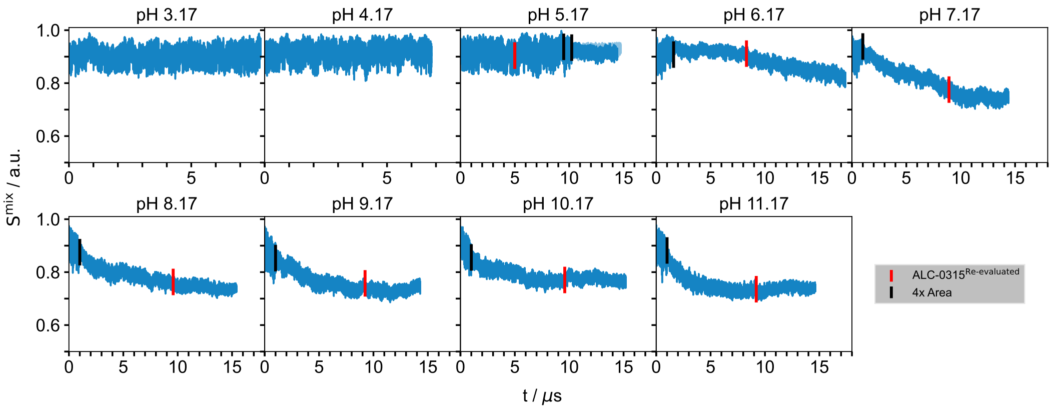


**Figure 7:** Conditional mixing entropy ($S^{mix}$) over time is shown for simulations of $D_{pH\ 5.17}$, $F^{1–6}$, $D^{‡}$, and $F^{0–6‡}$(see Tab. 1). $S^{mix}$ was calculated following the method of Brandani *et al.* (*1*), using a three-dimensional Euclidean distance cutoff of 1 nm to identify neighboring lipids. Lower values correspond to a stronger demixing of the lipid components. Black vertical lines indicate the quadrupling of the membrane surface, performed to prevent periodic boundary artifacts caused by an undersized surface area, while red vertical lines indicate the restart of the simulations with corrected polynomial coefficients (ALC-0315$^{\text{Re-evaluated}}$).

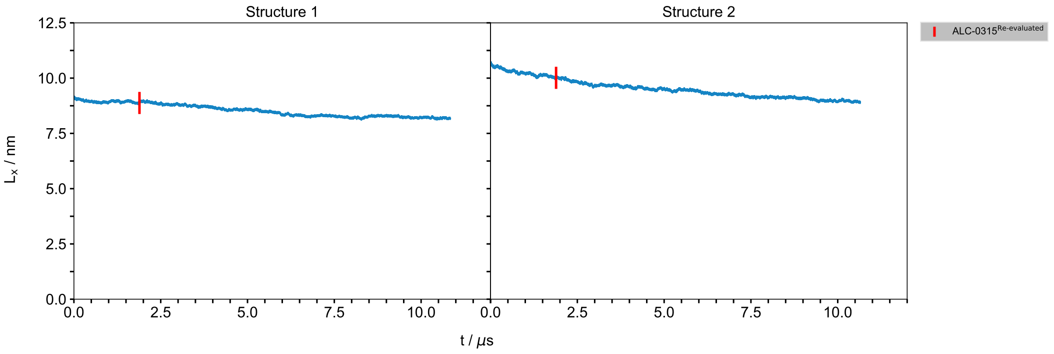


**Figure 8:** Length of the box vector in the x-direction over time for simulations of systems G and $G^{‡}$ (see Tab. 1) (LNP mixture containing mRNA). Black vertical lines indicate the quadrupling of the membrane surface, performed to prevent periodic boundary artifacts caused by an undersized surface area, while red vertical lines indicate the restart of the simulations with corrected polynomial coefficients (ALC-0315$^{\text{Re-evaluated}}$).

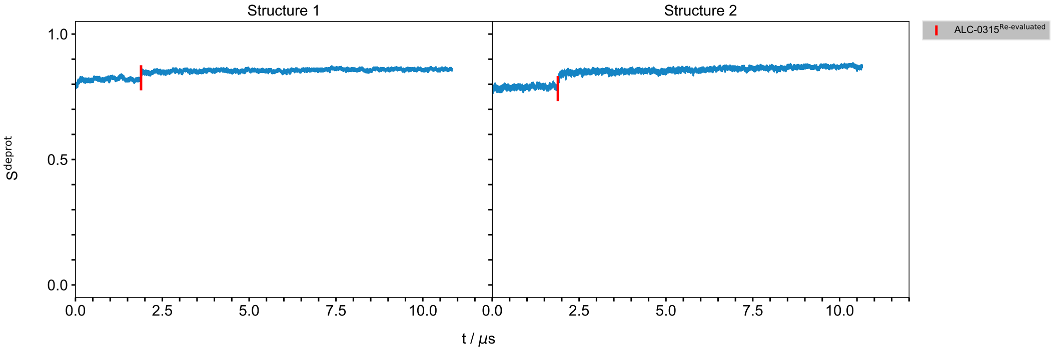


**Figure 9:** Fraction of deprotonation ($S^{deprot}$) (see Eq. 3) over time is shown for simulations of systems G and G$^{\ddagger}$ (see Tab. 1) (LNP mixture containing mRNA). The number of (de)protonated aminolipids was obtained via the $\lambda$-coordinate of each aminolipid ($\lambda < 0.2$, protonated; $\lambda > 0.8$, deprotonated) every 1 ps. Black vertical lines indicate the quadrupling of the membrane surface, performed to prevent periodic boundary artifacts caused by an undersized surface area, while red vertical lines indicate the restart of the simulations with corrected polynomial coefficients (ALC-0315$^{\text{Re-evaluated}}$).

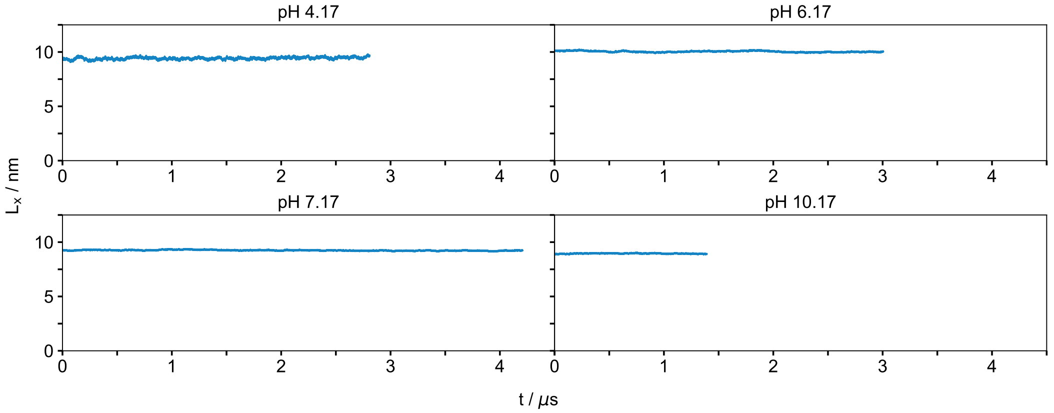


**Figure 10:** Length of the box vector in the x-direction over time for simulations of systems H$^{0-3\ddagger}$ (see Tab. 1) (LNP mixture containing TNS).

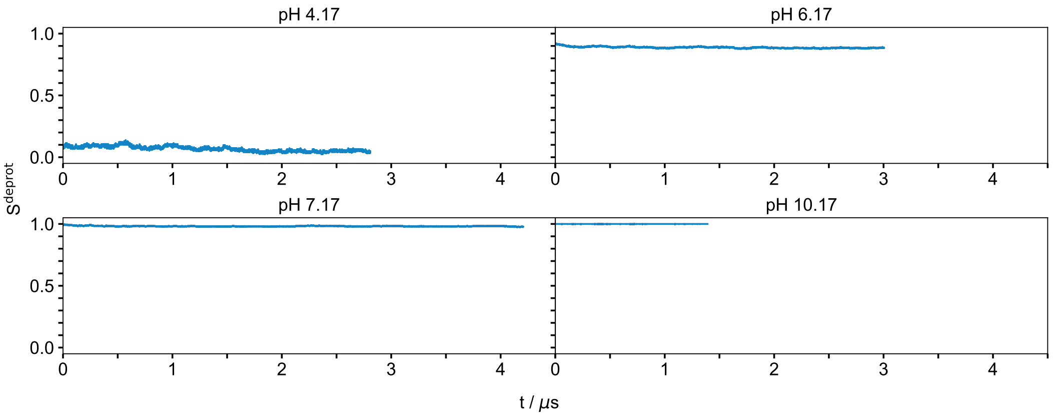


**Figure 11:** Fraction of deprotonation ($S^{deprot}$) (see Eq. 3) over time is shown for simulations of systems $H^{0–3‡}$ (see Tab. 1) (LNP mixture containing TNS). The number of (de)protonated aminolipids was obtained via the $\lambda$-coordinate of each aminolipid ($\lambda < 0.2$, protonated; $\lambda > 0.8$, deprotonated) every 1 ps.

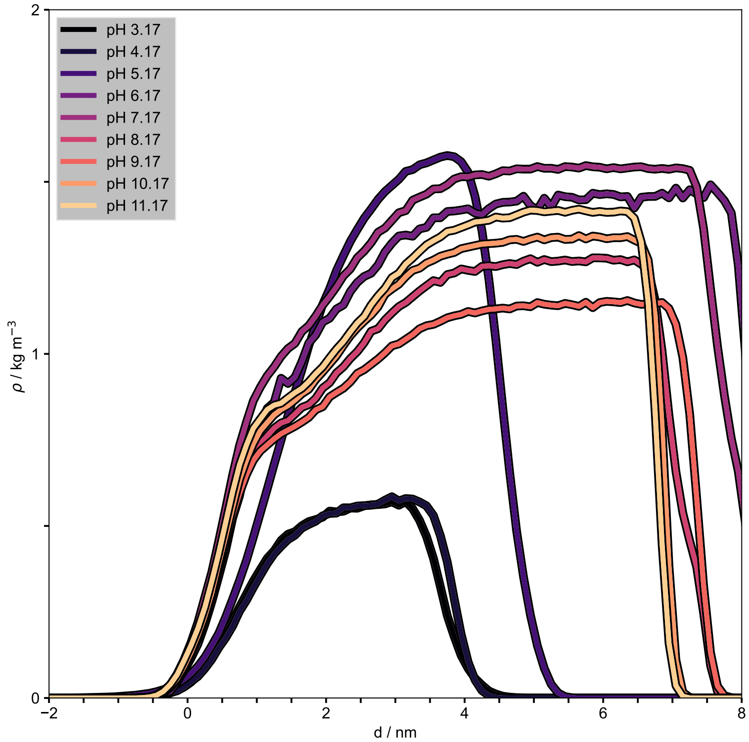


**Figure 12:** Average mass density profiles of the buffer particles relative to the membrane surface—defined by the median z-position of DSPC phosphorus atoms in a single leaflet—were calculated for systems $D^{‡}_{pH\ 3.17–4.17}$ and $F^{0–6‡}$(see Tab. 1). Averages were calculated after the $S^{deprot}$ values reached equilibrium; for details, see "Assessment of simulation convergence and error estimation" in the Methods section. As anticipated from the parameterization strategy of Buslaev *et al.* (*2*), the buffer particles avoid the hydrophobic region of the membrane.

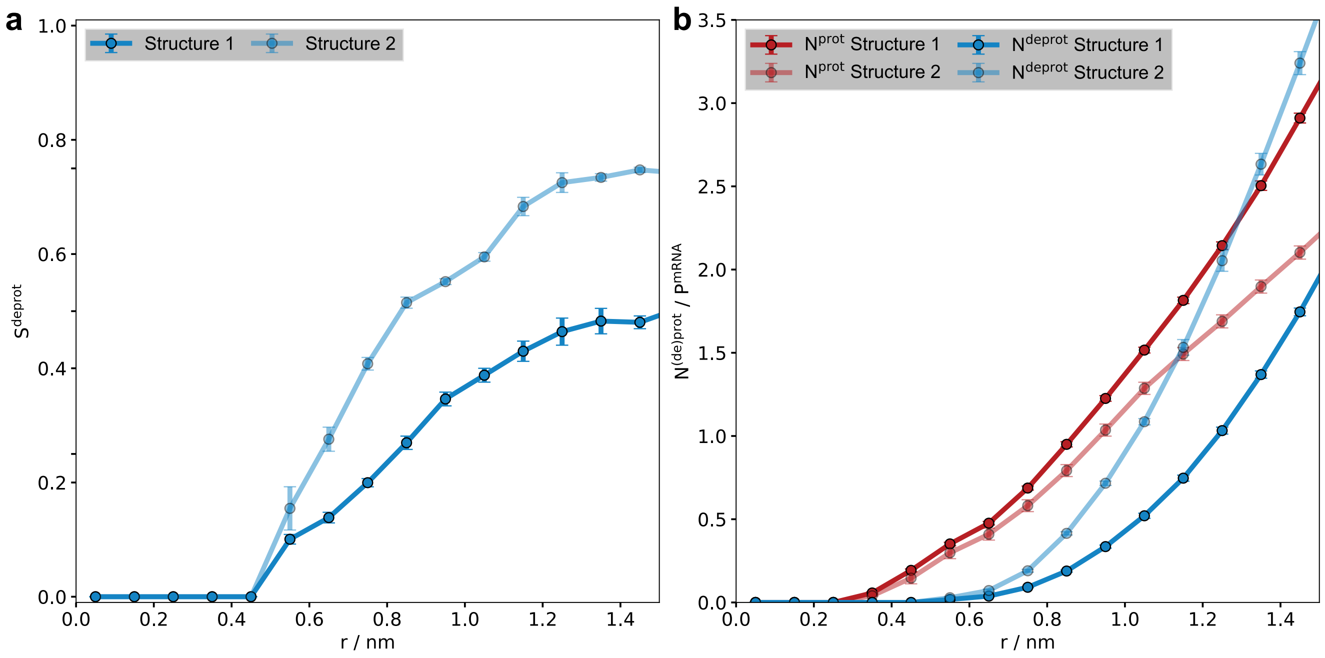


**Figure 13:** **a** Fraction of deprotonated ALC-0315 ($S^{deprot}$), **b** cumulative number of protonated ALC-0315 ($\lambda < 0.2$), and **c** cumulative number of deprotonated ALC-0315 ($\lambda > 0.8$) around the negatively charged backbone of the mRNA strands (see system G‡, Tab. 1). Note that the cumulative numbers are normalized by the amount of negatively charges phosphates in the mRNA backbone (here, $n^{PO_4^-}$=76). Distances were calculated between the nitrogen atom of ALC-0315 and the phosphorus atom in the nucleotide backbone. Averages were calculated after the $S^{deprot}$ values reached equilibrium.

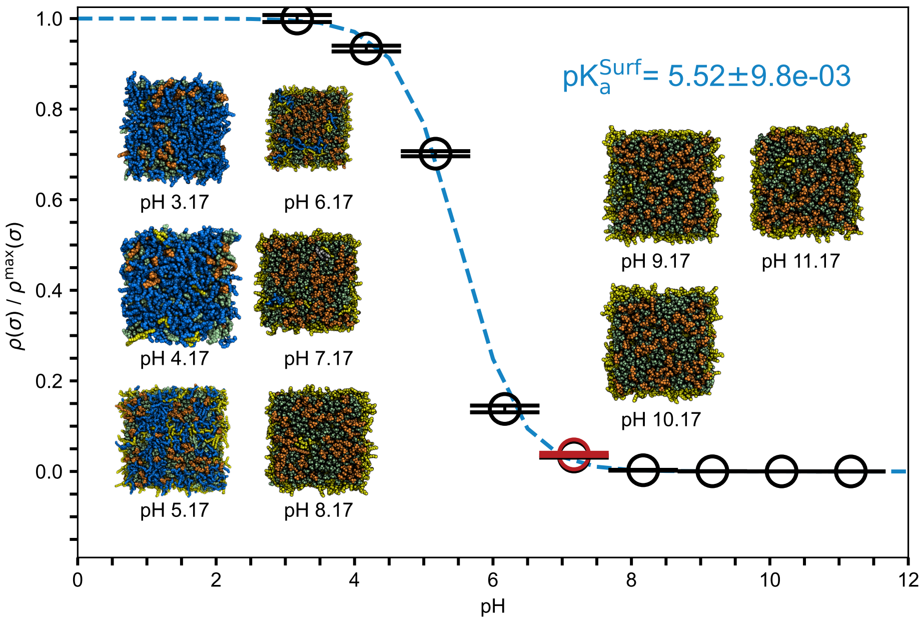


**Figure 14: pH-dependent surface charge density of LNP-mimetic systems.** The calculation included only protonated ALC-0315 aminolipids within 1.8 nm of, or above, the median position of the DSPC phosphorus atoms in one leaflet. The number of protonations was divided by the respective box area. Charge densities were then averaged over both leaflets in every frame. Error bars represent the standard error of the mean, calculated via block averaging (*3*). The values were normalized by the maximal charge density, $\rho^{max}(\sigma)$, derived from system D$^{\ddagger}$ at pH 3. The red marker represents the surface charge density of the LNP-mimetic system containing mRNA (Structure 1). The results indicate that the presence of mRNA inside the LNP does not affect the surface charge density at pH 7. Averages were calculated after the $S^{deprot}$ values reached equilibrium. Errors in the fit parameters were estimated using bootstrapping (*4*). Assuming that the scaled surface charge density is normally distributed at each pH value, new values were sampled from a normal distribution with an expectation value equal to the calculated average and a standard deviation equal to the standard error of the mean. The sampling was repeated 100,000 times; the reported pK$_a$ values corresponds to the mean of the bootstrap distribution, and the error bar represents its standard deviation. Inset images show the surface of one leaflet of the LNP-mimetic extracted from the last simulation frame: deprotonated ALC-0315 (*yellow*), protonated ALC-0315 (*blue*), DSPC (*orange*), and cholesterol (*green*). ALC-0159, and solvent are not shown. All pictures of atomistic structures were rendered with PyMOL (*5*).
